# PRMT5-mediated regulatory arginine methylation of RIPK3

**DOI:** 10.1101/2022.08.01.502351

**Authors:** Chanchal Chauhan, Ana Martinez Del Val, Rainer Niedenthal, Jesper Velgaard Olsen, Alexey Kotlyarov, Simon Bekker-Jensen, Matthias Gaestel, Manoj B. Menon

## Abstract

The TNF receptor-interacting protein kinases (RIPK)-1 and 3 are regulators of extrinsic cell death response pathways, where RIPK1 makes the cell-survival or death decisions by associating with distinct complexes mediating survival signaling, caspase activation or RIPK3-dependent necroptotic cell death in a context dependent manner. Using a mass spectrometry-based screen to find new components of the ripoptosome/necrosome, we discovered the protein-arginine methyltransferase (PRMT)-5 as a direct interaction partner of RIPK1. Interestingly, RIPK3 but not RIPK1 was then found a target of PRMT5-mediated symmetric arginine dimethylation. A conserved arginine residue in RIPK3 (R486 in human, R415 in mouse) was identified as the evolutionarily conserved target for PRMT5-mediated symmetric dimethylation and the mutations R486A and R486K in human RIPK3 almost completely abrogated its methylation. Rescue experiments using these non-methylatable mutants of RIPK3 demonstrated PRMT5-mediated RIPK3 methylation to act as an efficient mechanism of RIPK3-mediated feedback control on RIPK1 activity and function. Therefore, this study reveals PRMT5-mediated RIPK3 methylation as a novel modulator of RIPK1-dependent signaling.

## Introduction

TNF receptor-interacting protein kinases (RIPK)-1 and -3 are the master regulators of programmed cell death signaling. The concerted action of the RIP kinases downstream to the death receptors, interferon-alpha receptor (IFNαR) and Toll like receptor (TLR) regulate cell death/survival decisions during development and inflammation (Reviewed in (Humphries, Yang et al., 2015)). In addition, viral infection and genotoxic stress also activate RIPK1 and RIPK3. Recruitment to the receptor and subsequent ubiquitination of RIPK1 in the receptor associated complex I is crucial for the activation of MAP kinases and NFkB-dependent transcription, independent of RIPK1 kinase activity (Jiao, Wachsmuth et al., 2020, Kist, Komuves et al., 2021, Upton, Shubina et al., 2017, Zhang, Zhang et al., 2019). Activated RIPK1 associates with CASP8 and FADD to assemble a cytosolic death-inducing complex termed “ripoptosome” (complex IIb) resulting in apoptotic cell death (Micheau & Tschopp, 2003, Wang, Du et al., 2008). In the absence of caspase activity, active RIPK1 can also associate with RIPK3 via their RIP homotypic interaction motifs (RHIM) to form a functional hetero-amyloidal structure called necrosome (Cho, Challa et al., 2009, Li, McQuade et al., 2012). Subsequent RIPK3-mediated phosphorylation of MLKL leads to oligomerisation of MLKL followed by its translocation to the plasma membrane causing loss of membrane integrity and cell death defined as necroptosis. In addition to its canonical role as a pro-necroptotic kinase, RIPK3 also serves as an adaptor that drives CASP8-mediated apoptosis (Dondelinger, Aguileta et al., 2013, Mandal, Berger et al., 2014). Independent of the cell death events, RIPK3 also manifests pro-survival functions by regulating transcriptional responses (Moriwaki & Chan, 2017, Orozco, Daniels et al., 2019).

A whole lexicon of post-translational modifications regulates the pro-death and pro-survival functions of RIP kinases (Annibaldi & Meier, 2018, Delanghe, Dondelinger et al., 2020, Meng, Sandow et al., 2021). While RIPK1 phosphorylation by IKK1/2, MK2 and TBK1 were shown to act as checkpoints preventing RIPK1 activation and autophosphorylation at residue S166, mechanisms regulating the activity of RIPK3 are less understood (Dondelinger, Delanghe et al., 2017, Jaco, Annibaldi et al., 2017, Menon, Gropengiesser et al., 2017). Apart from being a substrate of RIPK1, RIPK3 also undergo auto-phosphorylation at residues T182, S199 and S227 (Choi, Park et al., 2018, McQuade, Cho et al., 2013, Sun, Wang et al., 2012). Members of the casein kinase 1 family phosphorylate RIPK3 at serine 227 and regulate its ability to recruit MLKL (Hanna-Addams, Liu et al., 2020). In addition, CHIP E3 ubiquitin ligase-mediated K48 ubiquitylation of RIPK3 at residues K55, K89, K363, and K501 and PELI1-mediated K48 ubiquitylation at K363 enhances RIPK3 turnover via proteasomal and lysosomal dependent degradation, respectively (Choi et al., 2018, Seo, Lee et al., 2016). Apart from this, O-GlcNAc transferase targets RIPK3 for O-linked GlcNAcylation and hinders necroptotic signaling (Li, Gong et al., 2019).

Protein methylation is known to influence various physiological processes including RNA processing, transcriptional regulation, DNA damage response, and cell cycle progression (Guccione & Richard, 2019, Martin & Zhang, 2005, Zhang, Wen et al., 2012). During the methylation reaction, a methyl group from S-adenosyl-L-methionine (SAM) is transferred to lysine or arginine residues in target proteins by highly specific methyltransferases. Protein arginine methyltransferases (PRMT) catalyse transfer of one or two methyl groups to arginine residues, giving rise to monomethyl-arginine, asymmetric dimethyl-arginine (aDMA) or symmetric dimethylarginine (sDMA). The family of mammalian PRMTs consists of nine members sorted into three groups: Type I methyltransferases (PRMT 1,3,4,6, and 8) generate aDMA by adding two methyl groups to the same terminal nitrogen of the arginine residue, Type II (PRMT5 and 9) mark each of the terminal nitrogen atoms of arginine with one methyl group, hence generating sDMA and the type III member PRMT7 catalyses mono-methylation of arginine residues (Reviewed in (Blanc & Richard, 2017)). Tight regulation of PRMTs is a prerequisite for regulated protein arginine methylation as there is no conclusive proof for the presence of specific arginine demethylases. PRMT activity is usually regulated by association with regulatory proteins, post-translational modifications of PRMTs as well as masking of their target sites by other modifications (Morales, Caceres et al., 2016). Interestingly, association of PRMT5 with WDR77/MEP50 is a prerequisite for PRMT5 activity in mammalian cells (Friesen, Wyce et al., 2002). Other interaction partners like RIOK1 and pICLN/CLNS1A are shown to regulate the substrate specificity by docking the methylosome complex to specific substrates (Guderian, Peter et al., 2011). PRMT5 activity and localisation is also regulated by its phosphorylation and methylation. While AKT-mediated T634 phosphorylation modulates membrane localisation of PRMT5 (Espejo, Gao et al., 2017), PKC_ι_-mediated S15 phosphorylation is a positive regulator of IL1-induced PRMT5 activity and NFκB-activation (Hartley, Wang et al., 2020a). Moreover, CARM1/PRMT4-mediated methylation of R505 residue is essential for the homodimerization and activation of PRMT5 (Nie, Wang et al., 2018).

Recent evidence indicate that the switch between pro-survival and pro-death functions of RIPK1 is regulated by a panel of modifications including phosphorylation and ubiquitination (Wang, Fan et al., 2021) and the presence of kinase checkpoints in RIPK1 activation revealed the presence of multiple stages in the maturation of the complex IIb/ripoptosome (Amin, Florez et al., 2018). To understand the assembly and maturation of the ripoptosome into an active death inducer, it is important to identify novel ripoptosome interactors. Through SILAC-based mass spectrometry (MS) analysis, we identified PRMT5 as a RIPK1 interaction partner. Subsequently, our studies reveal PRMT5-mediated symmetric di-methylation of RIPK3 as a novel mechanism regulating necrosome-mediated signaling.

## Results

### A screen for ripoptosome interactors by MS analysis identified PRMT5 as a RIPK1 interaction partner

To identify RIPK1 interactors during TNF-induced cell death, we first established a mouse-embryonic fibroblast (MEF) model system of induced expression of epitope-tagged RIPK1. *Ripk1^-/-^* MEFs were rescued by the expression of doxycycline (dox)-inducible FLAG-tagged RIPK1 and single clonal cell lines were isolated and stable labelled by isotopes. Dox-induced expression of RIPK1 was detectable in both clones analysed and RIPK1 is efficiently enriched by FLAG-tag based immune-precipitation. Immunoblots with anti-FLAG and anti-RIPK1 antibodies indicate that there is no detectable RIPK1 expression and enrichment in the non-induced cells (Dox -, Figure 1A). Since previous studies have shown that caspase inhibition can stabilize the complex IIb/ripoptosome (Tenev, Bianchi et al., 2011), we performed ripoptosome enrichment of cells treated with human TNFα in the presence of smac-mimetics (SM) and caspase inhibitor zVAD-fmk (zVAD). Efficient enrichment of ripoptosome-like complexes was verified by the stimulus-induced association of CASP8 and its cleaved form with FLAG-RIPK1 (Figure 1B). This enrichment is independent of RIPK1 phosphorylation by MK2, since it is detectable in the presence and absence of the MK2 inhibitor PF364402 (Fig. 1B). Immunoblot analyses revealed that both clones of rescued cells lacked detectable MLKL expression excluding the possibility of necroptotic cell death in this model (Supplementary Figure S1).

**Figure 1.**
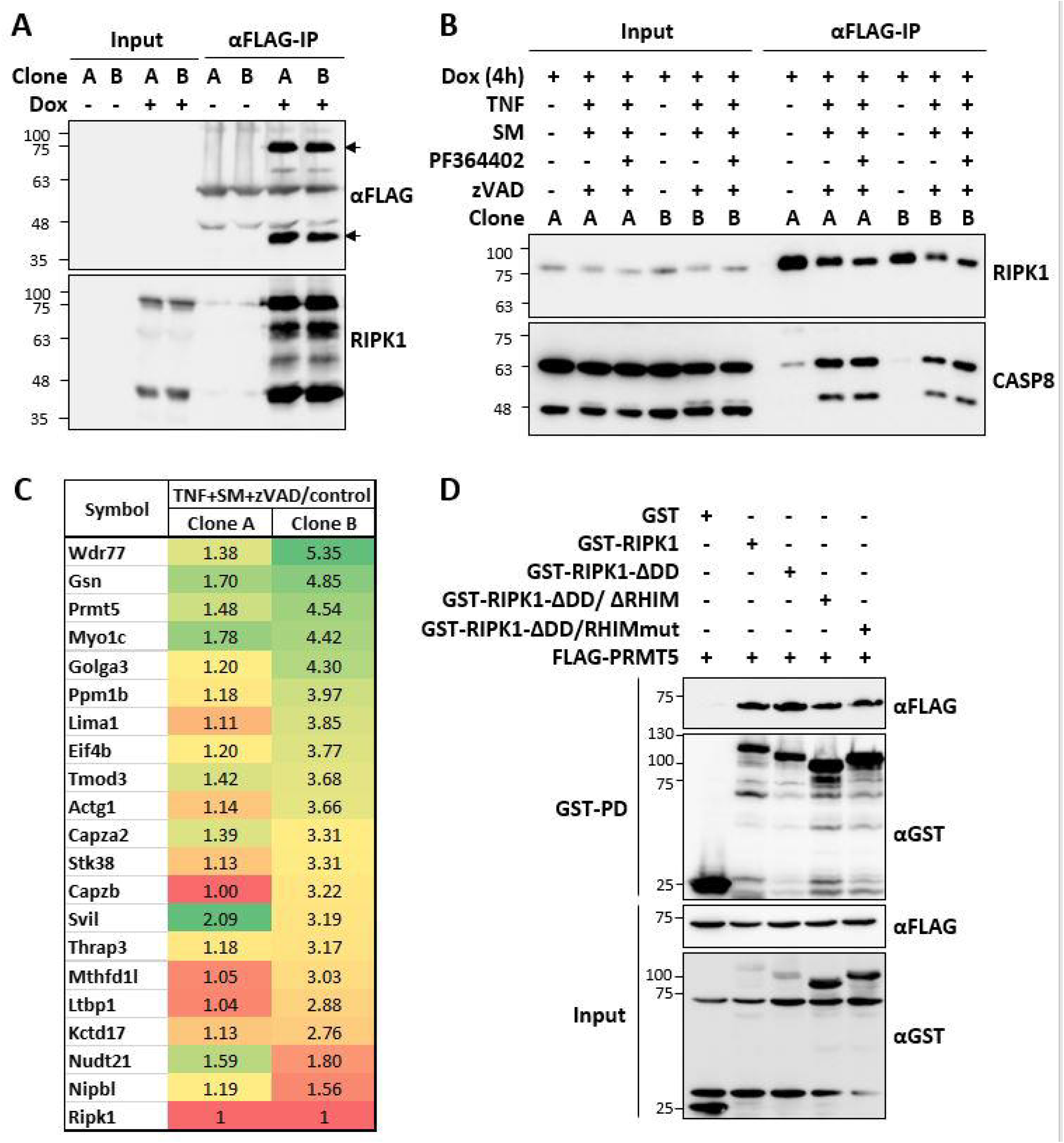
A screen for ripoptosome interactors identified PRMT5 as an interaction partner of RIPK1. **A**. Two clonal cell lines generated by retroviral transduction and single cell cloning of RIPK1-deficient fibroblasts with doxycycline (dox) inducible FLAG-tagged mRIPK1 expression vector were left untreated or treated with dox for 9 h. RIPK1 was enriched by FLAG-immunoprecipitation and detected in immunoblots with antibodies against FLAG tag and RIPK1. **B**. The cells were treated as indicated and subjected to FLAG-IPs as in panel A. Blots were probed with CASP8 antibodies to monitor assembly of the ripoptosome. **C**. Consistent RIPK1 interaction partners identified in both clonal lines by the mass spectrometry screen are represented with the fold enrichment between TNF+SM+zVAD treated and untreated conditions. **D**. GST-pulldown assays shows specific interaction of RIPK1 with PRMT5, independent of death domain (DD) and RHIM motif of RIPK1.

To identify ripoptosome interactors by a non-biased screen, we applied SILAC-based MS analysis. After the enrichment of FLAG-RIPK1 associated proteins from differentially SILAC-labelled control and TNF+SM+zVAD (Figure 1C) treated cells, the beads were pooled and eluted with FLAG peptides. The eluted proteins were digested and analyzed by LC-MS/MS in technical duplicates. The proteins which were identified in duplicate and consistent between the two independent clonal lines were considered as interactors (Figure 1C). In general, after normalization to enriched RIPK1 levels, clone B showed better enrichment for most identified proteins, but clone A also displayed significant enrichment. One of the consistent factors enriched was PRMT5, a type II methyl transferase. Interestingly, the PRMT5 methylosome consists of a hetero-octameric complex consisting of a central PRMT5 tetramer decorated with four WDR77 molecules. Remarkably, also WDR77 was amongst the most enriched proteins in our screen (Figure 1C) indicating enrichment of the entire PRTM5 methylosome. To verify the identified interaction between PRMT5 and RIPK1, we performed GST-pulldown experiments in HEK 293T cells transfected with FLAG-PRMT5 and GST-tagged mouse RIPK1 and mutants thereof. The results clearly showed strong enrichment of PRMT5 with GST-tagged full length mRIPK1 (1-656 aa) as well as C-terminal truncations of RIPK1 lacking the death domain (∆DD, mRIPK1-1-588 aa) or both death domain and RHIM motif (∆DD/∆RHIM, mRIPK1-1-500 aa) (Figure 1D). Similar results were obtained also when a dimerization deficient mutant of RIPK1 was used (∆DD, RHIMmut).

### RIPK3 is a target of PRMT5-mediated symmetric arginine dimethylation

Since the RIPK1 interactors included the core methylosome components PRMT5 and WDR77, which induces symmetric dimethylation of arginine residues, we monitored arginine methylation of RIPK1 after co-expression of PRMT5 by using a symmetric di-methyl arginine motif (SdmArg) antibody. Despite clear association of RIPK1 and PRMT5, there was no symmetric dimethylation detectable for GST-RIPK1 enriched using glutathione beads (Figure 2A). We then performed this analysis with RIPK3 and identified strong signals for symmetric dimethylation of RIPK3, which was enhanced upon PRMT5 co-expression in HEK 293T cells (Figure 2B). A strong interaction between RIPK3 and PRMT5 was also evident as GST-RIPK3 clearly co-purified co-expressed PRMT5 (Figure 2B). To verify that RIPK3 methylation is mediated by PRMT5, we monitored methylation in the presence and absence of the small molecule PRMT5 inhibitors GSK591 and LLY283. These PRMT5 inhibitors completely abrogated the methylation signals detectable by enrichment of GST-RIPK3 followed by immunoblotting with symmetric dimethylation specific antibodies (Figure 2C). Moreover, while the co-expression of wild-type PRMT5 enhanced the methylation of RIPK3, a catalytic deficient mutant of PRMT5 (PRMT5-R368A) failed to show any effect (Figure 2D). As a conclusive proof for PRMT5-mediated dimethylation of RIPK3, we performed siRNA experiments targeting endogenous PRMT5. Consistent with the previous findings, siRNA mediated depletion of PMRT5 completely inhibited the symmetric dimethylation of RIPK3 (Figure 2E). These findings support a model where RIPK1 and RIPK3 both interact with PRMT5, while only RIPK3 acts as a methylation target of PRMT5 (Figure 2F).

**Figure 2.**
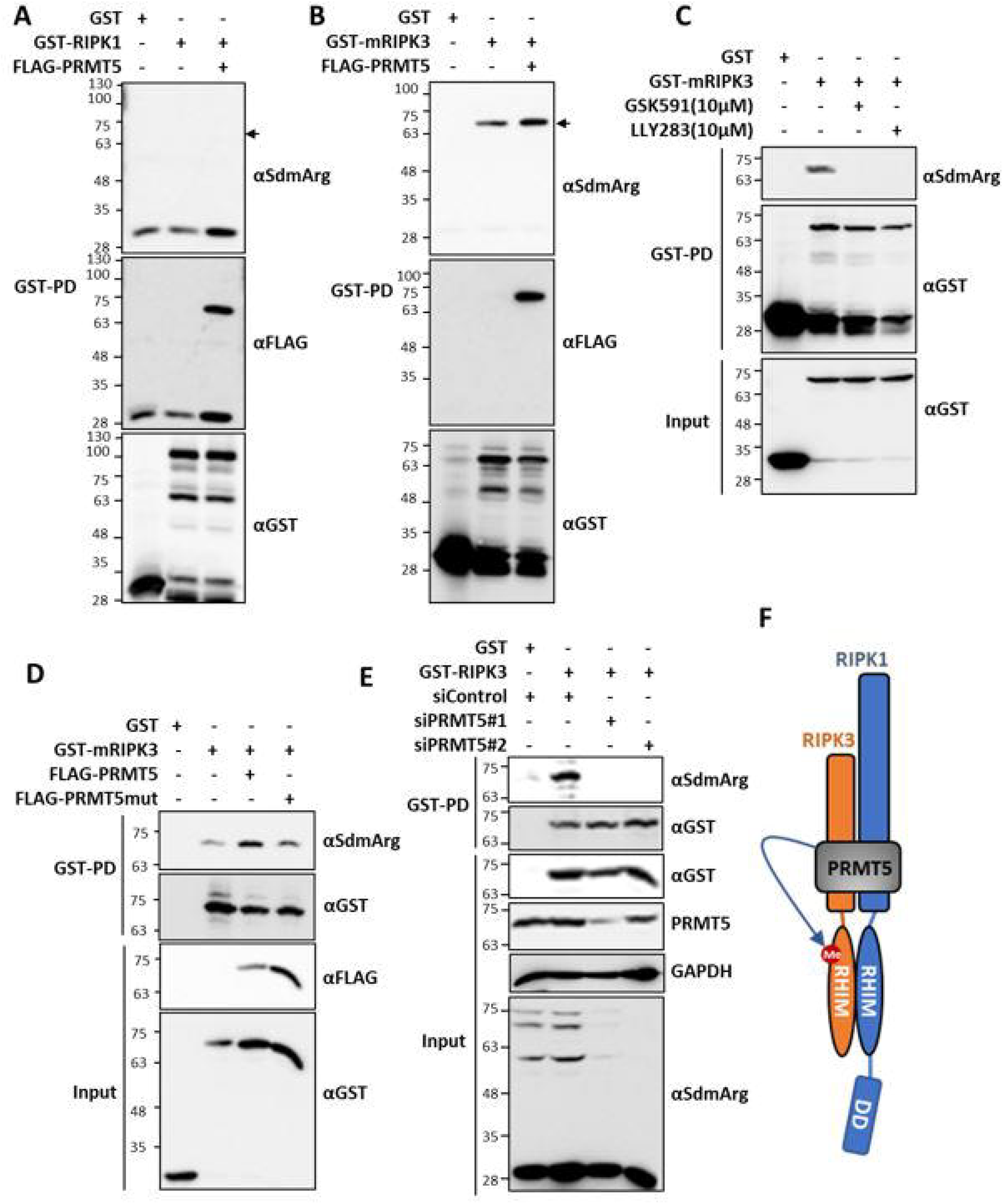
RIPK3 is a target of PRMT5-mediated symmetric arginine dimethylation. **A & B**. GST-enrichment and probing with antibodies against symmetric dimethyl arginine (αSdmArg) to monitor PRMT5-mediated dimethylation of mRIPK1 (A) and mRIPK3 (B). **C**. Effect of PRMT5 inhibitors GSK591 and LLY283 on RIPK3 methylation monitored by GST-enrichment and immunoblotting. **D**. FLAG-tagged wild-type PRMT5 but not the catalytic-deficient PRMT5 mutant (PRMT5mut, R368A) is capable of enhancing RIPK3 arginine methylation. **E**. siRNA mediated depletion of PRMT5 suppresses symmetric dimethylation of mRIPK3 in HEK293T cells. Efficient depletion of PRMT5 is shown by PRMT5 immunoblots and detection of methylated proteins in the total cell lysates **F**. While both RIPK1 and RIPK3 interacts with PRMT5, only RIPK3 is a substrate of symmetric arginine dimethylation.

### RIPK3 is methylated at a conserved arginine residue in the C-terminal tail

Despite the conservation of the cell death signaling pathways across species and high sequence similarity between human and mouse RIPK1, RIPK3 show more divergence between human and mouse (Figure 3A) with only 59% sequence identity (NP_006862.2 and NP_064339.2, respectively). Accordingly, RIPK3 and MLKL interactions display species specific preferences (Chen, Zhou et al., 2013) and major differences in the structural determinants of RIPK3-MLKL interaction and regulation between human and mouse proteins exist (Meng, Davies et al., 2021). To test whether human RIPK3 is also a target of PRMT5 and whether this modification is conserved across species, we monitored symmetric demethylation of hRIPK3 in the presence and absence of the two PRMT5-specific inhibitors GSK591 and LLY283. Human RIPK3 underwent strong arginine dimethylation in HEK 293T cells, which was completely lost upon PRMT5 inhibitor treatment (Figure 3B) or PRMT5 knockdown (Supplementary Figure S2). After demonstrating that the PRMT5-RIPK3 axis is conserved in humans, we looked at large-scale proteomic datasets for evidence for RIPK3 methylation. The PhosphoSite database (http://www.phosphosite.org) documents the existence of arginine methylation sites in both human and mouse RIPK3 outside the kinase domain (cf. Figure 3A). While the residues R264, R332, R413 and R415 are targets of methylation in mRIPK3, R422 and R486 are methylation sites on hRIPK3. Interestingly, R486 (R415 in mRIPK3) is the only conserved arginine methylation site between human and mouse RIPK3 (Figure 3C). We mutated this conserved arginine residue in human RIPK3 to alanine (R486A) or lysine (R486K) and monitored dimethylation of the mutants. Both mutations almost completely abrogated the symmetric dimethylation of RIPK3, establishing R486 as the predominant methylation site on RIPK3 (Figure 3D).

**Figure 3.**
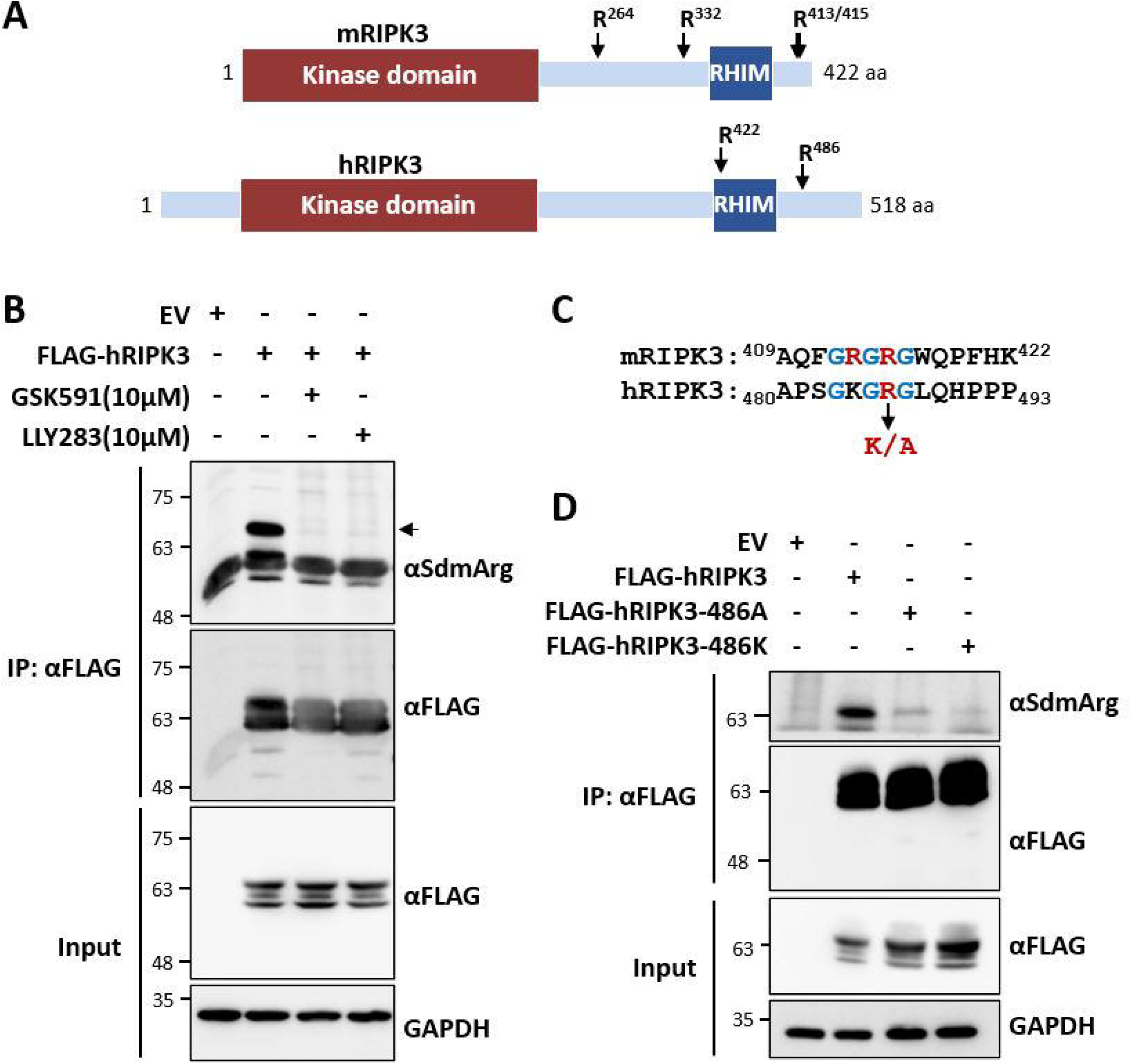
Identification of conserved arginine methylation sites on RIPK3. **A**. Schematic representation of human and mouse RIPK3 proteins with domain organisation and arginine (R) methylation sites. **B**. Human RIPK3 is a substrate of PRMT5-mediated symmetric arginine methylation. **C**. Alignment of mRIPK3 and hRIPK3 sequences reveal R486 in hRIPK3 (R415 in mRIPK3) as the only evolutionarily conserved methylation site. K/A denote mutagenesis of the site to lysine (K) and alanine (A) residues to generate methylation-deficient mutants. **D**. Mutagenesis of R486 residue (R486K or R486A) abrogates symmetric arginine dimethylation of hRIPK3 as shown by FLAG-IP enrichment and analysis with SdmArg antibodies.

### RIPK3 R486 modification provides feedback to RIPK1-RIPK3 signaling

To monitor RIPK3 functions and to investigate the significance of RIPK3 methylation, we established a rescue model of RIPK3 activity using the PANC1 human pancreatic cancer cell line. PANC1 cells do not express endogenous RIPK3 and, hence, cannot undergo necroptosis (Sun et al., 2012). We introduced RIPK3 into the PANC1 cells by lentiviral transduction and monitored necroptotic signaling (Figure 4A). RIPK3 expression was detected only in cells exogenously expressing RIPK3, while MLKL expression was visible in both cell lines. Upon treatment with the necroptotic stimuli (TNF+SM+zVAD), significant necroptosis-associated MLKL phosphorylation and strong downstream ERK1/2 activation were visible only in the RIPK3-rescued cells. Interestingly, RIPK1 autophosphorylation as indicated by the pS166 antibody signal was significantly suppressed only in the cells expressing RIPK3, indicating a RIPK3-mediated feedback control of RIPK1 autophosphorylation. A pre-treatment of the cells with the RIPK3 inhibitor GSK872 rescued RIPK1 autophosphorylation and suppressed necroptotic signaling, suggesting that the RIPK3-mediated feedback control on RIPK1 activation requires RIPK3 kinase activity. Moreover, inhibitors of oligomerization of both RIPK3 (PP2) and MLKL (NSA) also inhibited necroptotic signaling as well as RIPK3-mediated suppression of RIPK1-S166 phosphorylation (Figure 4A).

**Figure 4.**
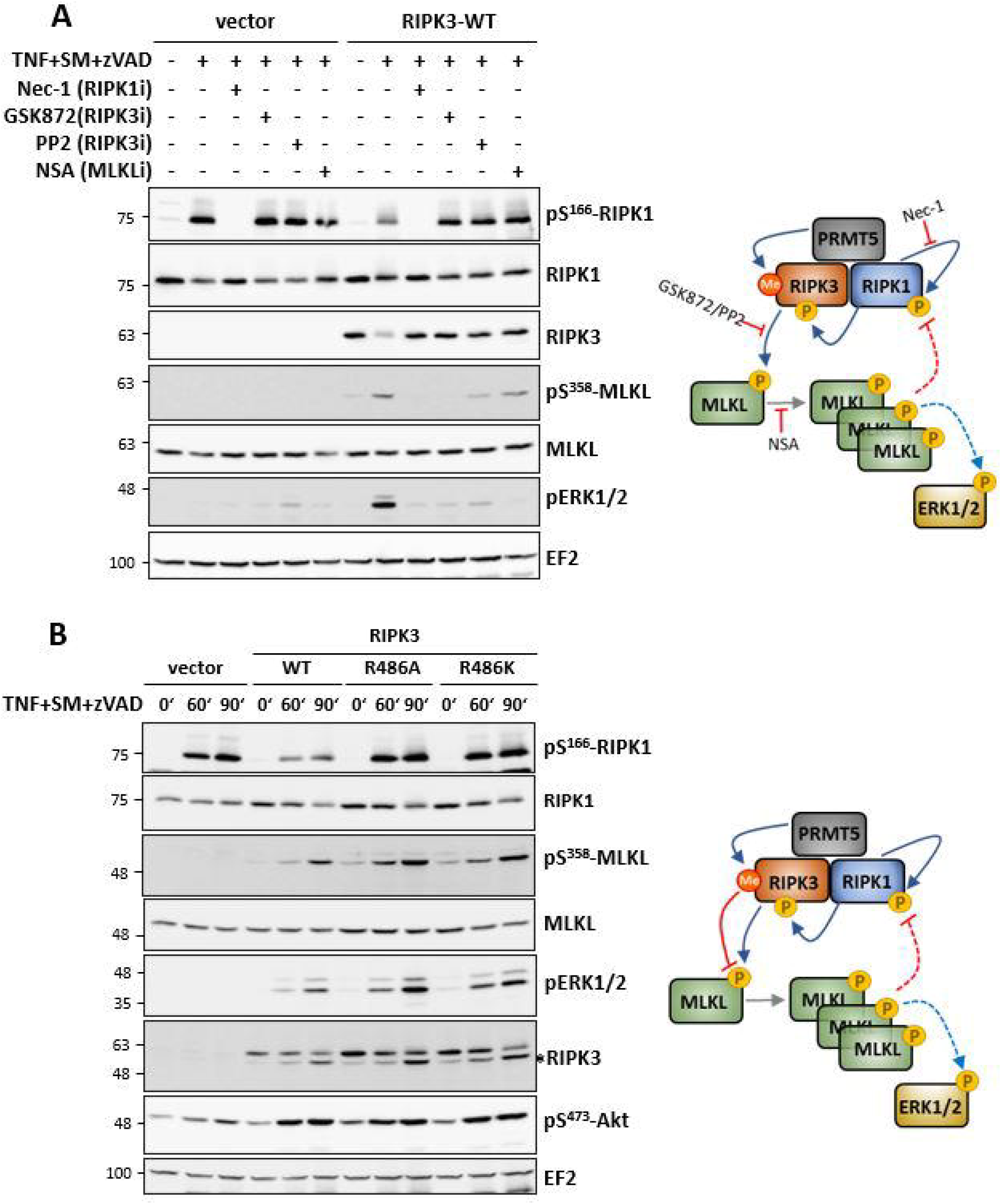
RIPK3 mediated feedback control on RIPK1 activity is dependent on RIPK3 methylation. **A**. The necroptosis-incompetent PANC1 cell line, which lacks RIPK3, was rescued by lentiviral transduction of FLAG-tagged wt-RIPK or empty vector. Necroptotic signaling was monitored by stimulating cells with TNFα, Smac-mimetics (SM) and zVAD-fmk for 90 minutes in the absence or presence of indicated kinase inhibitors (Nec-1, GSK872) or PP2 (RIPK3 oligomerisation inhibitor) or necrosulfonamide (NSA), an inhibitor of human MLKL oligomerisation. The right panel shows the schematic summary of the observations. RIPK3 expression facilitates necroptotic signaling in PANC1 cells (MLKL and ERK1/2 phosphorylation) but suppresses RIPK1 autophosphorylation (RIPK1-S166). The feedback control of RIPK3 on RIPK1 was abrogated by RIPK3 and MLKL inhibitors. **B**. PANC1 rescue model as in panel A comparing the necroptotic signaling in cells transduced with WT and methylation site mutants (R486K and R486A) of FLAG-tagged RIPK3. The corresponding panel to the right summarises the conclusions from the immunoblots. Top right panel depicts the signaling effects and the processes targeted by the inhibitors used. RIPK3 activation by RIPK1 leads to MLKL phosphorylation and oligomerisation, with downstream ERK1/2 signaling. The oligomerisation of MLKL also seems necessarr for suppressing pS166-RIPK1 phosphorylation. The lower panel scheme depicts the possible effect of RIPK3 methylation in the same system. RIPK3-mediated suppression of RIPK1-S166 is lost upon methylation site mutation. In addition, the necroptotic signaling is also enhanced in the absence of R486 methylation. This indicate an inhibitory role for RIPK3-methylation in RIPK3 downstream signaling including the feebback control of RIPK1 activity.

To further investigate the role of PRMT5-mediated methylation of RIPK3 at residue R486, we generated PANC1 cells rescued with wild-type (WT) and methylation-deficient mutants (R486K or R486A) of RIPK3. After transduction and selection of cell lines for uniform transduction, the selected empty vector and RIPK3 expression lines showed similar expression levels of GFP expressed from the bidirectional promotor demonstrating uniform transduction and similar expression levels of WT and mutant RIPK3 expression (Supplementary Figure S3). When necroptotic signaling was compared between the four cell lines, we observed clear suppression of RIPK1 autophosphorylation by WT-RIPK3 again, however, the R486A and R486K mutant cell lines consistently displayed a stronger signal for pS166-RIPK1 similar to those observed in RIPK3-deficient empty vector transduced cells (Figure 4B). The necroptotic downstream signaling indicated by MLKL phosphorylation and ERK1/2 activation was enhanced in the cells expressing the K486 mutated RIPK3, indicating that methylation acts as negative effector of overall necroptotic signaling. However, phosphorylation of Akt/PKB in response to necroptotic stimulus was enhanced in RIPK3 or mutant expressing cells, obviously independent of RIPK3 methylation (Figure 4B). In contrast to the negative effect of RIPK3 on necroptotic RIPK1 autophosphorylation, S166 phosphorylation of RIPK1 in response to a pro-apoptotic stimulus (TNF+SM) was not suppressed by RIPK3 expression. Interestingly, in this case a dose dependent increase of S166 phosphorylation was observed (Supplementary Figure S4).

## Discussion

The ripoptosome/complex IIb formed during death receptor-dependent and -independent cell death consists of the core components RIPK1, FADD and CASP8. Inhibition of CASP8 activity leads to the association of RIPK3 with this complex, leading to the formation of a RIPK1-RIPK3 containing necrosome capable of phosphorylating MLKL and inducing necroptosis. In the last years, many interaction partners and posttranscriptional modifications of RIPK1 and RIPK3 were identified as modulators of these cell death pathways. In the present study, we provide the first proof for an arginine dimethylation of RIPK3 as such a modulatory mechanism. Using mass spectrometry approaches, we have identified the protein arginine methyl transferase PRMT5 as a RIPK1 interactor which mediates a symmetric arginine dimethylation of RIPK3 C-terminus. Using a RIPK3 rescue model in a RIPK3-deficient pancreatic cancer cell line which normally cannot undergo necroptosis, we demonstrate that the evolutionary conserved dimethylated arginine residue R486 is important for RIPK3-mediated feedback control on RIPK1 autophosphorylation. Hence, we established a link between PRMT5 and the necroptosis regulator RIPK3. A recent study also corroborates our finding by showing that PRMT5 exhibits anti-tumor effects via regulation of necroptosis (Otani, Sur et al., 2021). PRMT5-depletion was shown to sensitise glioblastoma cells to the antitumor effects of the protein phosphatase 2A inhibitor LB100 by facilitating necroptosis.

Interestingly, the ripoptosome interactome detected in our MS screen included several methylosome components including RIOK1 and WDR77 in addition to PRMT5 (Fig. 1 and Supplementary Table 1). This indicates a potentially more general role for protein arginine methylation in the regulation and/or maturation of the ripoptosome. It is important to note that one of the earliest reports on PRMT5 function was its interaction with death receptors and consequential inhibition of apoptosis by promoting NFkB activation (Tanaka, Hoshikawa et al., 2009). This study also revealed that while PRMT5 interacted with the TRAIL receptors, there was no association with the TNFR1 and PRMT5 regulated TRAIL sensitivity in cancer cells. However, more recent evidence indicates a role for PRMT5 downstream to TNF receptors in mounting an NFkB-dependent inflammatory response. PRMT5 has been shown to enhance NFkB activation by methyating R30 and R174 residues on NFkB p65 subunit (Harris, Chandrasekharan et al., 2016, Wei, Wang et al., 2013). PRMT5-mediated YBX1-R205 methylation was also shown to affect NFkB-dependent gene expression promoting colon cancer cell proliferation and anchorage-independent growth (Hartley, Wang et al., 2020b). CARM1/PRMT4-mediated methylation of PRMT5 was shown to be essential for its homodimerisation and activation leading to histone methylation and suppression of gene expression (Nie et al., 2018), while S15 phosphorylation of PRMT5 by PKCι acts as a postive modulator of PRMT5 activity and PRMT5-dependent NFκB activation (Hartley et al., 2020a). Interestingly, PRMT4 is known to regulate a subset of NFκB-target genes by CBP300 methylation downstream to TNF and TLR4 signaling (Covic, Hassa et al., 2005). CFLAR (CASP8 and FADD-like apoptosis regulator), also known as c-FLIP is another TNF receptor downstream molecule which is a target of arginine methylation (Li, An et al., 2019). A role for PRMTs including PRMT5 in inflammatory gene expression is gaining prominence and incidentally we also observed a role for PRMT5-mediated RIPK3 methylation in necrosome downstream MAPK activation. The methylation-deficient mutants consistently gave enhanced RIPK3-dependent ERK1/2 phosphorylation signals in response to necroptotic stimulus (Figure 4). RIPK1 and RIPK3 mediated inflammatory response if mediated by ERK1/2 kinases, independent of the cell death (Najjar, Saleh et al., 2016).

We also observed a detectable increase in the methylation-deficient RIPK3 mutant protein levels compared to the wild-type RIPK3 in the rescued PANC1 cells. The reason for such an effect is not clear. However, despite moderately elevated levels, RIPK3 mutants were not effective in providing the feedback suppression on RIPK1 autophosphorylation. Ubiquitination and subsequent lysosomal degradation of RIPK3 by the CHIP-E3 ubiquitin ligase was shown as a mechanism of necroptosis suppression (Seo et al., 2016). Interestingly, PRMT5 is also a proven substrate of CHIP mediated ubiquitination and proteasomal degradation (Zhang, Zeng et al., 2016). In lung cancer patients, an expression signature with low CHIP expression and high RIPK3 expression is associated with bad prognosis (Kim, Chung et al., 2020). Regulation of RIPK3 levels could indeed be a mechanism relevant to inflammation and cancer.

The mutual regulation of RIPK1 and RIPK3 is still poorly understood. RIPK1 is the upstream activator of RIPK3. However, it is also known as a factor preventing spontaneous RIPK3 oligomerisation and activation (Orozco, Yatim et al., 2014). While RIPK1-mediated necroptosis, but not CASP8-dependent apoptosis, requires RIPK3 and MLKL, artificial oligomerization of RIPK3 can result in both apoptosis and necroptosis. In the absence of RIPK1, oligomerised RIPK3 predominantly induces MLKL-dependent necroptosis, but also apoptosis in MLKL-deficient cells (Cook, Moujalled et al., 2014). Studies using genetic models of RIPK1-deficiency has shown a kinase activity-independent scaffolding role for RIPK1 in preventing epithelial cell apoptosis and necroptosis (Dannappel, Vlantis et al., 2014). Cells from the RIPK3-RHIM mutant mice were protected from necroptosis as well as apoptosis further confirming the complex signaling interplay of these protein kinases in the regulation of different pathways of cell death (Zhang, Wu et al., 2020). This study also revealed the presence of enhanced RIPK1 autophosphorylation and RIPK1-dependent lymphoproliferative disease in the RIPK3-RHIM mutant mouse in a *Fadd*^-/-^ background suggesting a role for RIPK3-mediated feedback on RIPK1 mediated inflammatory response. PRMT5-mediated methylation of RIPK3 may participate in the modulation of this feedback. Further studies using genetic mouse models will be necessary to reveal the complex interplay of PRMT5 and RIPK3 in inflammation and to identify additional substrates of PRMT5-mediated methylation in the RIPK1/3 RHIM interactome.

## Materials and Methods

### Cell Culture

3T3-immortalised *Ripk1-/-* MEFs were kindly provided by Prof. K. Ruckdeschel, (Universitätsklinikum Hamburg-Eppendorf, Germany). *Ripk1-/-*MEFs, HEK293T cells and PANC-1 cells were cultured in DMEM supplemented with 10% heat-inactivated fetal calf serum (FCS) and 1% Penicillin/Streptomycin. The cell lines were maintained at 37⍰°C and with 5% CO2 in a humidified atmosphere.

### Antibodies and Reagents

MLKL (#14993), αSdmArg (#13222), RIP3 (#13526), Caspase-8 (#4927), pS166-RIPK1 (#65746), pS358-MLKL (#91689), pS345-MLKL (#37333), phospho-p38 MAPK (Thr180/Tyr182) (#9211), phospho-p44/42 MAPK (Erk1/2) (Thr202/Tyr204) (#4370) and pS473-Akt (#4060) antibodies were from Cell Signaling Technology (CST). Further antibodies used were against GST (sc-138, Santa Cruz Biotechnology), RIPK1 (#610459, BD Biosciences), GAPDH (#MAB374, Millipore), EF2 (sc-166415, Santa Cruz Biotechnology), FLAG (#F3165, Sigma-Aldrich) and GFP (sc-9996, Santa Cruz Biotechnology). The secondary antibodies used were anti-rabbit IgG-HRP (Conformation Specific) (#5127, CST), mouse TrueBlot^®^ ULTRA (#18-8817-33, Rockland Immunochemicals), goat anti-mouse IgG (H+L)-HRP (#115-035-003, Dianova) and goat anti-rabbit IgG (H+L)-HRP (111-035-003, Dianova).

The following reagents at given concentrations were used for cell treatments: Doxycycline (D9891, Sigma, 1µg/ml), recombinant human TNFα (rHuTNF, #50435.50, Biomol, 10⍰ng/ml), Birinapant/Smac Mimetics (HY-16591, MedChem Express, 1µM), pan-caspase inhibitor zVAD-fmk (4026865.0005, Bachem, 25⍰μM), PF3644022 (4279, Tocris, 5μM), GSK591 (Cay18354-1, Cayman, 10µM), LLY-283 (HY-107777, MedChem Express, 10µM), Nec-1 (BML-AP309-0020, Enzo Life Sciences, 50µM), PP2 (HY-13805, MedChem Express, 10µM), NSA (5025, Tocris, 10µM), GSK872 (HY-101872, MedChem Express, 5µM).

### Plasmids, cloning and mutagenesis

Dox-inducible pSERS retroviral vector as described previously (Menon, Sawada et al., 2014) was converted to a Gateway Destination vector and a FLAG-tagged mRIPK1 cDNA (Menon et al., 2017) was shuttled in to create doxycycline-inducible retroviral expression system for low level RIPK1 expression. The human and mouse RIPK3 coding region (NM_006871.4 and NM_019955.2, respectively) PCR-amplified from HeLa cell and MEF cell cDNA, respectively, were cloned into the pENTR-D-TOPO directional cloning vector. L-R clonase II mediated shuttling was used to generate N-terminally tagged expression vectors. 3xFLAG-PRMT5 and 3xFLAG-PRMT5-R368A mutant expression vectors were described previously (Bruns, Grothe et al., 2009). The expression vector pCR3.V62-Met-FLAG-RIP3 was reported earlier (Menon et al., 2017). Site-directed mutagenesis was performed using the QuikChange mutagenesis kit (Agilent) to generate FLAG-RIP3-486A and FLAG-RIP3-486K methylation site mutants. pLBID-MCS-GFP-P2A-Puro was gifted by Dr. A. Schambach (MHH, Germany). pLBID-FLAG-hRIPK3 were generated by subcloning PCR products from the pCR3.V62-Met vector into the AgeI/XhoI sites. All other plasmids were described previously (Menon et al., 2017). All cloning and mutagenesis primer sequences used are listed in Supplementary Table S2.

### Transient transfection

HEK293T cells were transfected using polyethylenimine (PEI; Sigma-Aldrich). Transfected cells were maintained in antibiotic free DMEM media supplemented with 10% FCS for 12–16⍰hours followed by providing with complete DMEM media. Transfected cells were analysed between 20 and 36⍰hours post transfection.

### Retroviral transduction

Ripk1-/-MEFs were stably rescued with Doxycycline inducible retroviral vectors expressing mouse RIPK1. For preparation of viral supernatants, 7.5 million Ecopack-HEK293T cells were seeded in 10 cm plates and transfected overnight with 5 µg each of pCL-Eco and pSERS11 based retroviral vectors using PEI, in antibiotic free medium with 10% FCS. Medium was changed to fresh complete media supplemented with 1X non-essential amino acids (NEAA). Virus containing supernatants were harvested five times from transfected Ecopack-HEK293 cells (Clontech/Takara), over a period of three days post-transfection and filtered with 0.45 µm filters. 1.2x 105 Ripk1-/-MEF cells were seeded per well in a 6 well plate and were cultured for 4 days with viral supernatants in the presence of 8µg/ml polybrene. RIPK1 expression was monitored by intracellular staining and flow cytometry analysis using an Accuri-C6 cytometer (BD Biosciences).

### Lentiviral transduction

To generate a rescue model for RIPK3, PANC-1 cells (RIPK3-deficient) were transduced with lentiviral vectors expressing wt/mutant human RIPK3 or pLBID-MCS-GFP-P2A-Puro empty vector. ViraPower Lentiviral Expression System (Invitrogen) was used to package pLBID lentiviral vectors expressing FLAG-hRIPK3 or methylation site mutants or empty vector. Viral supernatants were collected 48 h post transfection and filtered through 0,45µm filters as described previously (Menon et al., 2017). PANC-1 cells were transduced twice with the supernatant containing lentivirus and 8µg/ml polyberene followed by positive selection of transduced cells in presence of 1-2µg/ml puromycin. Flow cytometry analysis of GFP was used to monitor comparable transduction efficiency between the cell lines.

### Intracellular Flow cytometry staining of RIPK1

Cells were suspended in PBS at a concentration of 10^7^ cells/ml and were fixed with 3-volumes of 4% PFA. After 30 minutes at room temperature, fixed cells were washed and permeabilized for 30 minutes with ice-cold 90% methanol. After 2x washes with PBS, cells were blocked with 4% BSA on ice for 30 minutes. After 1h at RT with anti-RIPK1 antibody (BD Biosciences # 610459, 1:100 diluted in 1% BSA-PBS) cells were incubated with 1:700 diluted secondary antibodies (anti-mouse Alexa Fluor-488, Invitrogen) for 30 minutes. Samples were washed with PBS and analysed with Flow cytometry using an Accuri-C6 cytometer (BD Biosciences).

### SILAC-based Mass Spectrometry analysis of RIPK1 interactome

Post transduction and sorting, two clonal cell lines were generated from dox-inducible FLAG-RIPK1 rescued *Ripk1^-/-^* MEFs. These clones were metabolically labelled with light (Lys0, Arg 0) and medium (Lys4, Arg 6) non-radioactive isotope amino acids. SILAC Protein Quantitation Kit (Trypsin), DMEM (A33972, ThermoFisher Scientific) was used for generating light labelled cell lines and for generation of medium labelled cell lines, L-Lysine-2HCL,4,4,5,5-D4 (ThermoFisher Scientific) and 13C-labelled L-Arginine HCl (201203902, Silantes) were individually purchased. Cells were cultured in respective medium supplemented with 200 µg/ml of L-Proline for at least 10 passages for achieving incorporation of each respective labels. Cells were seeded at a density of 3×10^6^ cells in 10 cm plates and were treated the next day with doxycycline (1µg/ml) for 4 h, followed by treatment of light labelled cells with only DMSO control and of medium labelled cells with TNF+SM+zVAD for 2hours. FLAG-RIPK1 bound complexes were enriched from 1.5mg of protein lysate per sample. Pre-cleared lysates were incubated with 40µl of 50% ANTI-FLAG^®^ M2 affinity gel (#A2220, Sigma) on a rotor for 3 hours at 4°C. After immunoprecipitation, Anti-FLAG beads were pooled from the two differentially treated light and medium labelled samples for each clonal cell line. Proteins were eluted from pooled beads with 0.5mg/ml FLAG peptide (#F3290, Millipore Sigma) by shaking at 1000 rpm for 1 hour at 4°C. Samples were separated on a 10% SDS-PAGE gel and in-gel digested overnight with Trypsin/Lys-C protease mix and resulting peptides were desalted using C18-stage tip. The purified peptides from each sample were analysed by mass spectrometry in technical duplicates. Samples were analyzed on the Evosep One system using an in-house packed 15 cm, 150 μm i.d. capillary column with 1.9 μm Reprosil-Pur C18 beads (Dr. Maisch, Ammerbuch, Germany) using the pre-programmed gradients for 60 samples per day (SPD). The column temperature was maintained at 60°C using an integrated column oven (PRSO-V1, Sonation, Biberach, Germany) and interfaced online with the Orbitrap Exploris 480 MS. Spray voltage was set to 2.0 kV, funnel RF level at 40, and heated capillary temperature at 275°C. Full MS resolutions were set to 60,000 at m/z 200 and full MS AGC target was 300 with an IT of 22 ms. Mass range was set to 350−1400. Full MS scan was followed by top 10 DDA scans, using 1.3m/z isolation window, 45,000 resolution, 86 ms injection time, AGC target of 200 and normalized collision energy was set at 30%. All data were acquired in profile mode using positive polarity and peptide match was set to off, and isotope exclusion was on. The intensity of each peptide was normalised with intensity of RIPK1 in respective light and medium labelled samples. SILAC ratios were then calculated by comparing intensities of medium labelled peptides to intensities of light labelled peptides (Supplementary Table S1). Proteins with SILAC ratios more than 1 were considered and compared for consistency between technical replicates and clonal cell lines.

### siRNA mediated knockdown of PRMT5

Two siRNAs targeting human PRMT5 (FlexiTube siRNA Hs_PRMT5_1, cat#SI04216492 and Hs_PRMT5_2, cat# SI04248951, Qiagen) were used for knockdown experiments in HEK 293T cells. The siRNA sequences are provided in the Supplementary Table S2. 5×10^5^ cells were seeded in 6 cm plates and transfected with siRNAs using Lipofectamine™ RNAiMAX Transfection Reagent (#13778030, Invitrogen) as per manufacturer protocol. Transfected cells were cultured for 48 hours. Cells were then re-transfected overnight with mRIPK3 and hRIPK3 expression vectors. The next day, fresh media was provided to the transfected cells and plates were washed and frozen 24h post transfection.

### Western Blotting

Cells were lysed directly in cell culture plates with 2x SDS sample buffer containing 10% SDS, 1.5M TRIS pH 8.8, 5% Glycerol, 2.5% 2-Mercaptoethanol, and Bromophenol Blue. The samples were then boiled for 5 minutes at 95°C. Denatured protein lysates were run on SDS-PAGE (7.5–16% gradient) gels and immunoblotted to Nitrocellulose blotting membrane (#10600001, Amersham Biosciences). Western blots were blocked with 5% powdered skim milk in PBS with 0.1% Tween 20 for 1h at room temperature. After three washes with PBS containing 0.1% Tween 20, membranes were incubated overnight with the primary antibody at 4 °C followed by 1 hour incubation with horseradish peroxidase-conjugated secondary antibodies at room temperature. Blots were developed with an ECL detection kit, WESTAR NOVA 2.0 (#XLS07105007, 7Biosciences), and the digital chemiluminescence images were taken by a Luminescent Image Analyser LAS-3000 (Fujifilm). Bands were quantified using ImageJ software.

### Immunoprecipitation of FLAG-tagged proteins and GST-pulldowns

GST pulldowns and FLAG-IPs were performed as described previously (Menon et al., 2017, Ronkina, Lafera et al., 2016). Cells were lysed in lysis buffer containing 20mM Tris-acetate pH 7.0, 0.1mM EDTA, 1mM EGTA, 1mM Na3VO4, 10mM b-glycerophosphate, 50mM Sodium Fluoride, 5mM Na-pyrophosphate, 1% Triton X-100, 1mM benzamidine, 2µg/ml leupeptin, 0.1% 2-Mercaptoethanol, 0.27M Sucrose, 0.2mM PMSF, 1µg/ml Pepstatin A, 1X Protease Inhibitor Cocktail (B14002, Bimake), Phosphatase Inhibitor Cocktail (B15001, Bimake). Lysates were then cleared by centrifugation and the beads were blocked with 1% BSA to avoid non-specific binding of PRMT5 to Glutathione beads. For the enrichment of the fusion proteins, pre-cleared lysates were incubated with Monoclonal ANTI-FLAG^®^ M2 affinity gel (A2220, Sigma) or Protino Glutathione Agarose 4B (745500.10, Macherey-Nagel) for 16 hours at 4°C. The beads were then washed 4X with immunoprecipitation wash buffer containing 1X TBS, 50mM Sodium fluoride, 1% Triton X-100, 1mM Na_3_VO_4_. The beads were boiled in 2x SDS sample buffer for 5 minutes at 95°C and the eluted protein complexes were analysed by Western blotting.

### Statistics and Reproducibility

All Immunoblot results presented are representative results from at least three independent experiments.

## Acknowledgements

We thank Dr. Klaus Ruckdeschel (UKE-Hamburg) for the gift of RIPK1 KO Mouse-embryonic fibroblasts, Dr. Peter Claus (Hannover Medical School, Germany) for the gift of PRMT5 expression vectors, Dr. Axel Schambach (Hannover Medical School, Germany) for sharing the pLBid lentiviral expression vector and Dr. Sven Diederichs (DKFZ-Heidelberg) for providing PANC1 cells. This work was supported by the Deutsche Forschungsgemeinschaft (DFG) grants ME4319/3-1 and GA453/16-1. Work at The Novo Nordisk Foundation Center for Protein Research (CPR) is funded in part by a generous donation from the Novo Nordisk Foundation (Grant number NNF14CC0001). MS-based proteomics work was also funded by grant EPIC-XS-823839.

