## Supplementary Figure S1 for "PRMT5-mediated regulatory arginine methylation of RIPK3"

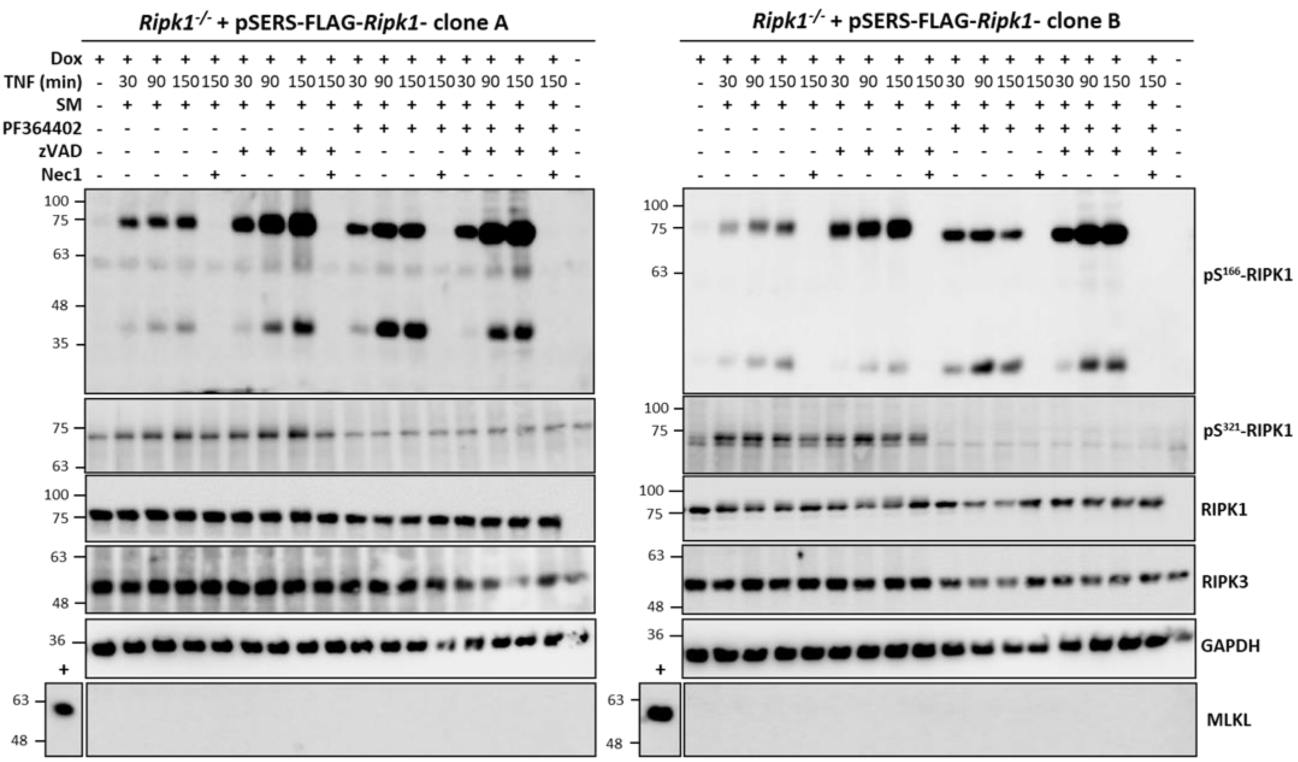


**Supplementary Figure S1. No MLKL expression during cell death signaling in the cell lines used for MS analyses.** Two independent rescued *Ripk1*-KO cell lines (Clone A and B) were treated as indicated and the cell lysates were analysed with antibodies against pS166-RIPK1, pS321-RIPK1, RIPK1, RIPK3, GAPDH and MLKL. MLKL expression was not detected in these cells indicating lack of necroptosis. A positive control sample (MK2$^{-/-}$ mouse embryonic fibroblast lysates) was loaded to show that MLKL antibody was functional.
