## Supplementary Figure S2 for "PRMT5-mediated regulatory arginine methylation of RIPK3"

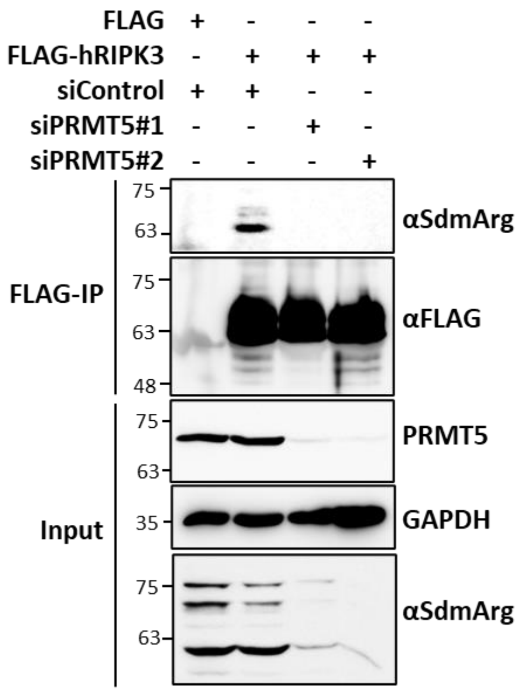


**Supplementary Figure S2. PRMT5 knockdown abrogated hRIPK3 methylation.** FLAG-tagged human RIPK3 was immunoprecipitated from HEK293T cell transfected with siRNAs targeted against PRMT5 or control siRNA. The samples were probed with anti-symmetric dimethyl arginine antibodies. Control blots show efficient knockdown of PRMT5 and effective downregulation of general methylation signals in the input cell lysates.
