## Supplementary Figure S3 for "PRMT5-mediated regulatory arginine methylation of RIPK3"

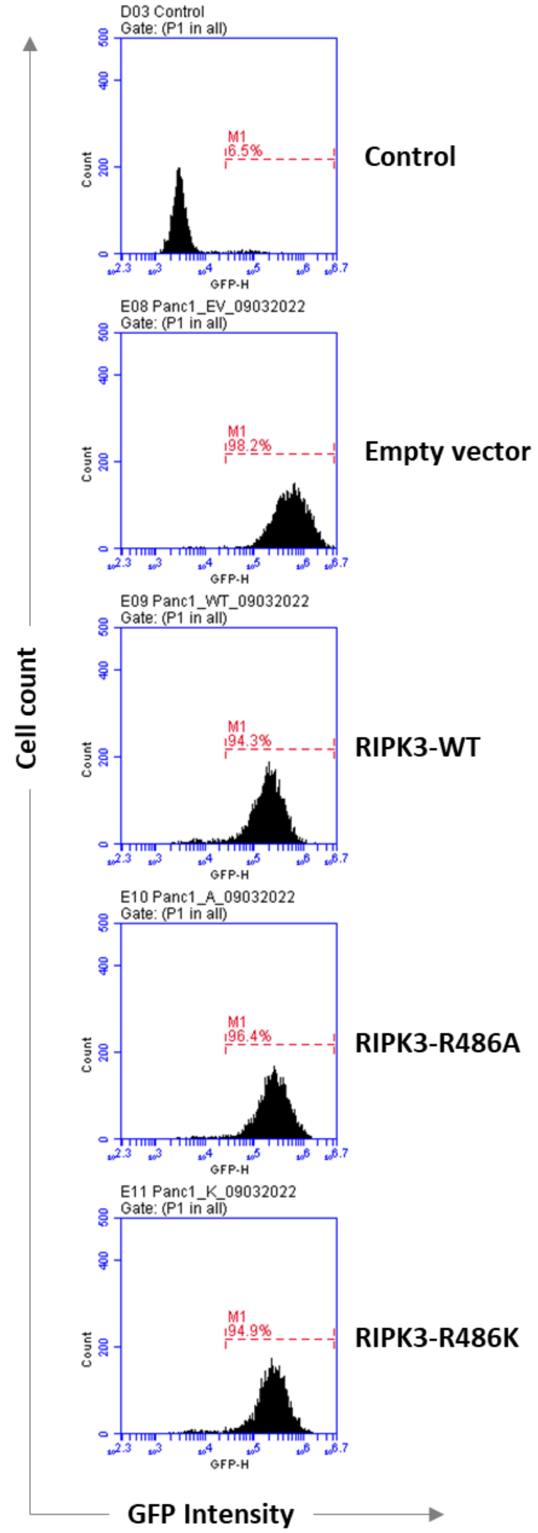


**Supplementary Figure S3. Flow-cytometry analyses of RIPK3-rescued PANC1 cells.** PANC1 cells transduced with the pLBID-GFP backbone vectors were selected by culturing in the presence of puromycin and the GFP levels analyzed by flow-cytometry. WT and mutant RIPK3 expressing cells showed comparable expression levels and transduction efficiency.
