## Supplementary Figure S4 for "PRMT5-mediated regulatory arginine methylation of RIPK3"

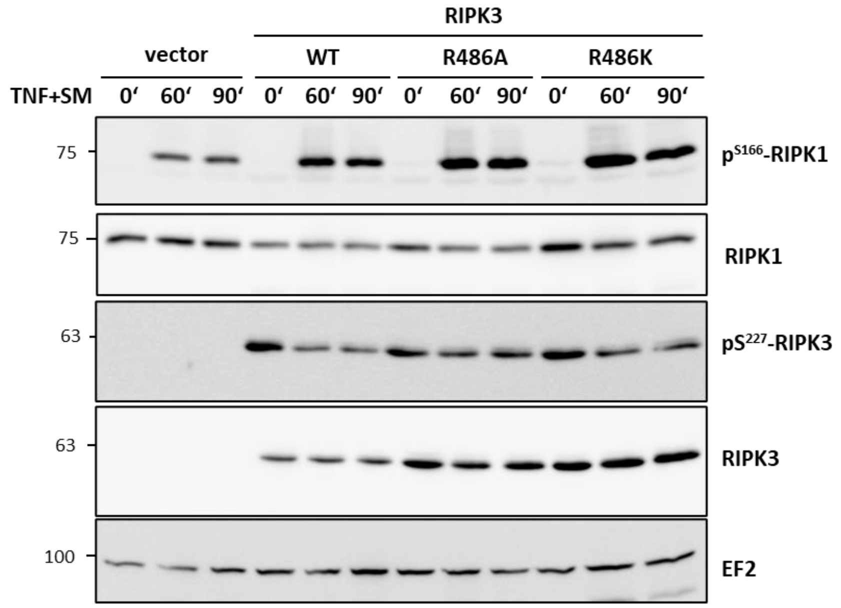


**Supplementary Figure S4. Role of RIPK3 in RIPK1 autophosphorylation in response to pro-apoptotic stimulus.** PANC1 rescue model as in Figure 4A comparing the apoptotic signaling in cells transduced with Wild-type and methylation site mutants (R386K and R386A) of FLAG-tagged human RIPK3. The cell lysates were probed with indicated antibodies after Immunoblotting to detect RIPK1 and RIPK3 activation and levels. EF2 is shown as sample-loading control.
